# Nanoluciferase-based Method for Detecting Gene Expression in *C. elegans*

**DOI:** 10.1101/745927

**Authors:** Ivana Sfarcic, Theresa Bui, Erin C. Daniels, Emily R. Troemel

## Abstract

Genetic reporters such as the green fluorescent protein (GFP) can facilitate measurement of promoter activity and gene expression. However, GFP and other fluorophores have limited sensitivity, particularly in tissues that autofluoresce like the intestine of the nematode *Caenorhabditis elegans*. Here, we present a highly sensitive Nanoluciferase (NanoLuc)-based method in multi-well format to detect constitutive and inducible gene expression in *C. elegans*. We optimize detection of bioluminescent signal from NanoLuc in *C. elegans* and show that it can be detected at 400,000-fold over background in a population of 100 animals expressing intestinal NanoLuc driven by the *vha-6* promoter. We can reliably detect signal in single *vha-6p::Nanoluc-*expressing worms from all developmental stages. Furthermore, we can detect signal from 1/100 dilution of lysate from a single *vha-6p::Nanoluc*-expressing adult and from a single *vha-6p::Nanoluc*-expressing adult “hidden” in a pool of 5,000 N2 wild-type animals. We also optimized various steps of this protocol, which involves a lysis step that can be performed in minutes. As a proof of concept, we used NanoLuc to monitor promoter activity of the *pals-5* stress/immune reporter and we were able to measure 300 and 50-fold increased NanoLuc activity after proteasome blockade and infection with microsporidia, respectively. Altogether, these results indicate that NanoLuc provides a highly sensitive genetic reporter for rapidly monitoring gene expression in *C. elegans*.

## Introduction

Genetically encoded reporters are important tools to monitor gene expression, and provide faster read-outs than other methods. For example, promoter-driven reporters are often used as proxies for assessing mRNA expression, and are faster than more direct measurements of mRNA expression, such as quantitative reverse transcriptase polymerase chain reaction (qRT-PCR) or single-molecule fluorescence in situ hybridization (smFISH). In *C. elegans*, genetic reporters have traditionally been introduced into the genome as multi-copy arrays that contain hundreds to thousands of copies (Mello et al. 1991). While multiple copies of a transgene can increase reporter signal, a small RNA-mediated process called transgene silencing often reduces expression from these multi-copy reporters in *C. elegans* (De-Souza et al. 2019; Minkina and Hunter 2018). Because of this phenomenon, measurement of reporter gene expression in *C. elegans* can be confounded by factors that regulate transgene silencing instead of regulating expression of the gene of interest itself. Generation of low-copy array strains (Schweinsberg and Grant 2013) or integration of reporters as single copy transgenes via MosSCI (Mos-1 mediated single-copy insertion) or CRISPR/Cas9 (Dokshin et al. 2018; Frøkjaer-Jensen et al. 2008) can reduce or eliminate silencing, but has the disadvantage of producing lower signal than multi-copy transgenes (Mendenhall et al. 2015). The most commonly used genetically encoded reporters include fluorophores like the green fluorescent protein (GFP). Unfortunately, weak background fluorophores present in multicellular organisms can decrease the overall sensitivity of such fluorophore-based assays. In particular, gut granules of the *C. elegans* intestine are highly auto-fluorescent and can hamper measurement of gene expression from this tissue (Teuscher and Ewald 2018). Thus, there is a need for genetically encoded reporters in *C. elegans* with better overall signal than fluorophores like GFP.

In contrast to fluorescent reporters, bioluminescent reporters generate *de novo* light without the need for external excitation through photons, and they are highly sensitive with a broad dynamic range (Thorne, Inglese, and Auld 2010). Bioluminescent signal is generated through oxidation of a substrate (Luciferin) by a Luciferase enzyme and there are many Luciferin/Luciferase pairs. These reporters are commonly used in mammalian systems and less often in *C. elegans*. To date, the ATP-dependent firefly luciferase has been used in *C. elegans* to measure mitochondrial function, larval molting, feeding behavior and circadian rhythm, and also to monitor viral infection (Gammon et al. 2017; Lagido, McLaggan, and Glover 2015; Olmedo et al. 2015; Palikaras and Tavernarakis 2016). While the ATP dependence of firefly luciferase is a means to monitor mitochondrial activity, the requirement for ATP hampers the ability of this luciferase to accurately investigate other processes like gene expression, because a lack of signal may simply reflect lowered ATP levels in the cell (Brock 2012). Therefore, the use of highly sensitive, ATP-independent luciferases would facilitate broader use of bioluminescent reporters in *C. elegans*.

In this study we establish the ATP-independent Nanoluciferase (NanoLuc) as both a constitutive and inducible genetically encoded luciferase reporter in *C. elegans*. NanoLuc was developed by Promega (Hall et al. 2012), who optimized a subunit of the *Oplophorus gracilirostris* deep sea shrimp luciferase to generate a reporter with small size, high physical stability, and high brightness. Promega further recommends to use the coelenterazine 2-furanylmethyl-deoxy-coelenterazine (Furimazine) as a substrate for NanoLuc to achieve a high luminescent signal with increased half-life and decreased autoluminescence compared to other luciferase/luciferin pairs. The 19 kDa monomer NanoLuc exerts an ATP-independent glow-type blue signal (Emission max 460 nm) with a half-life longer than two hours. Compared to firefly and renilla luciferases, NanoLuc emits a ∼150-times brighter signal, has better assay stability at higher temperatures, and operates at a wider pH range and in the presence of urea (Hall et al. 2012). NanoLuc has been used for several applications in non-*C. elegans* systems, like mammalian cells (England, Ehlerding, and Cai 2016). Here we used the MosSCI technique to generate single-copy reporter strains that have constitutive and inducible expression of NanoLuc in *C. elegans*. We used these strains to develop a sensitive and quantitative plate-based assay that has the ability to detect bioluminescent signal in a fraction of a worm constitutively expressing NanoLuc, and to detect induction of NanoLuc being driven by an immune/stress-regulated promoter.

## Materials and Methods

### *C. elegans* Strains and Culture

N2 was used as the wild-type strain. Transgenic reporter strains ERT412, ERT513, ERT529 and ERT729 are described in Table 1. All transgenic strains carry single-copy insertions on chromosome II introduced by the MosSCI method (Frøkjaer-Jensen et al. 2008) and were backcrossed at least three times before use. All *C. elegans* strains were maintained at 20°C on Nematode Growth Media (NGM) plates seeded with *E. coli* OP50-1 bacteria according to standard methods (Brenner 1974). Stocks of synchronized, starved first larval stage (L1) animals were generated by bleaching gravid adults (Emmons, Klass, and Hirsh 1979).

**Table 1.** Transgenic Reporter Strains used in this study.

| Strain | Genotype |
| --- | --- |
| ERT412 | *unc-119(ed3) III; jySi20[vha-6p::strepII::3XFLAG::GFP::let-858 3' UTR; cb-unc-119(+)] II* |
| ERT513 | *unc-119(ed3) III; jySi35[vha-6p::Nanoluc::unc-54 3' UTR; unc-119 (+)] II* |
| ERT529 | *unc-119(ed3) III; jySi40[vha-6p::Nanoluc::3XFLAG::unc-54 3' UTR; unc-119 (+)] II* |
| ERT729 | *unc-119(ed3) III; jySi44[pals-5p::Nanoluc::unc-54 3' UTR; unc-119 (+)] II* |

### Fluorescence Microscopy

Worms were anesthetized with 10 mM levamisole in M9 buffer and mounted on 5% agarose pads for imaging. Images were captured with a Zeiss Axio.Imager M1 using similar exposure times for Differential Interference Contrast (DIC) and GFP measurements for both images in Figure 1A and B.

**Figure 1.**
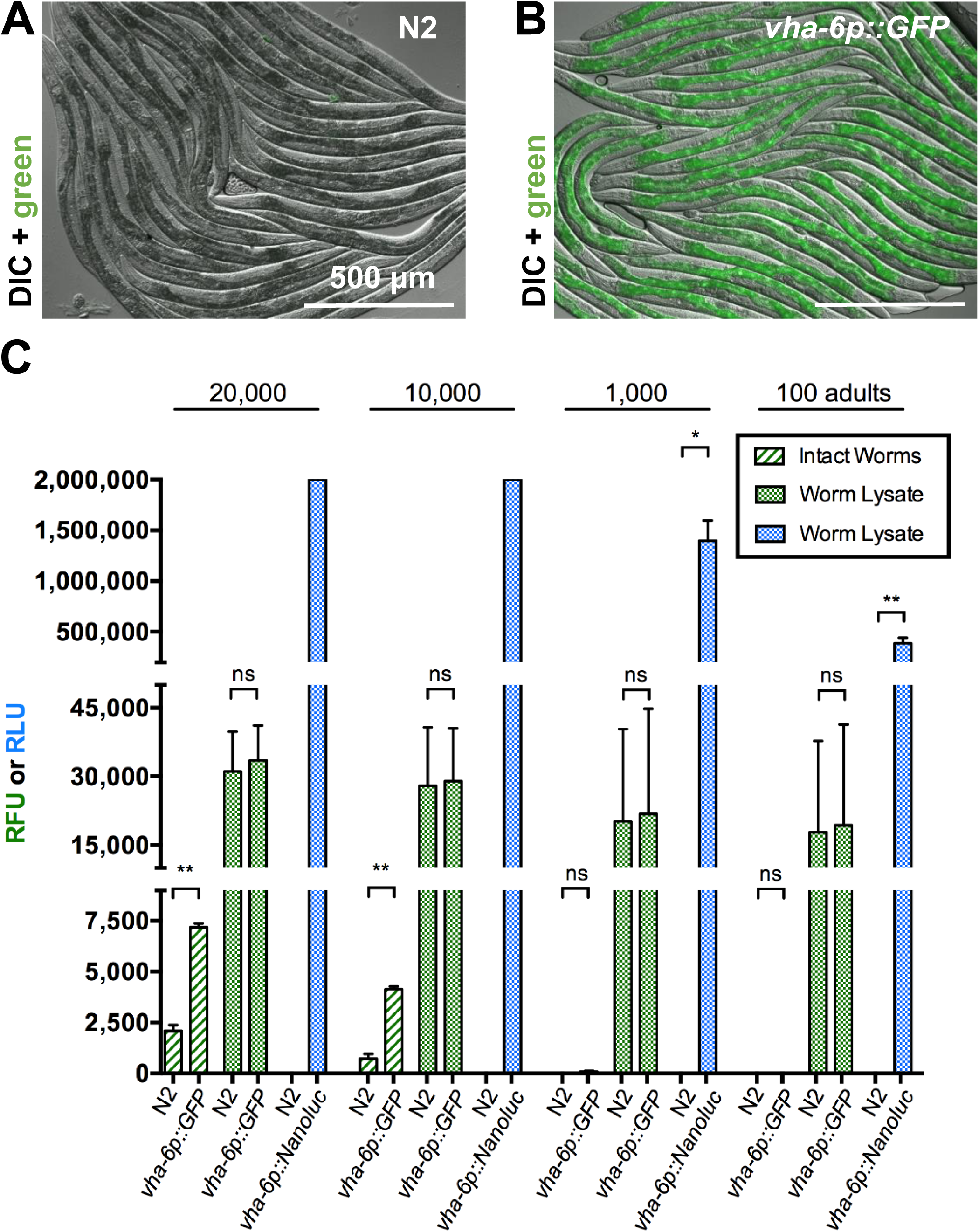
Nanoluciferase (NanoLuc) Sensitivity Exceeds GFP Sensitivity on Plate Reader. (A, B) Fluorescence microscopy images show strong intestinal expression of GFP in *vha-6p:: GFP* transgenic *C. elegans* but not in N2 adults. Images obtained by Differential Interference Contrast (DIC) microscopy with GFP fluorescence overlay. (C) Fluorescent Signal (Relative Fluorescent Units, RFU, green) or Luminescent Signal (Relative Luminescent Units, RLU, blue) measured on a plate reader from intact young adult worms and worm lysate of either N2, *vha-6p::GFP* or *vha-6p::Nanoluc* animals. RFU were measured with 485 nm Excitation and 520 nm Emission filters using 10 flashes per well and cycle. RLU were measured for 1 second, 10 minutes after addition of Furimazine (NanoGlo Reagent, Promega) without filters. n=2-4 trials for each condition. Error bars are Standard Deviation (SD), ** *p* < 0.01, * *p* < 0.05, ns = not significant.

### GFP Measurement on Plate Reader

For both N2 and ERT412, synchronized L1s were grown to the adult stage at 20°C and GFP was measured in either intact animals or in lysate from disrupted animals. In both cases, animals were first washed with M9 with 0.1% Tween 20 (M9-T) to remove excess OP50-1. For measurement of intact animals, 100, 1,000, 10,000 or 20,000 young adults in 200 μl M9-T were transferred to a black 96-well assay plate with clear bottom. To prepare worm lysate, 100, 1,000, 10,000 or 20,000 young adults in 210 μl M9-T with protease inhibitor (Roche, cOmplete mini, EDTA-free) were vortexed on a Disruptor Genie vortexer (Scientific Industries) in microfuge tubes with 15-20 Silicon Carbide beads (BioSpec Products, #11079110sc, 1 mm diameter) for 5 minutes at 4°C. These vortexed samples were spun at 20,000xg for 5 minutes, and then 200 μl of the worm lysate was transferred to a black, clear-bottom 96-well assay plate (Costar, #3603). Fluorescent signal in intact and lysed worms was measured on a NOVOstar plate reader (BMG Labtech), detecting 10 flashes per well and cycle using a 485 nm Excitation and a 520 nm Emission filter. M9-T was used as a blank for measurements of intact animals, and M9-T vortexed with Silicon Carbide beads was used as a blank for measurements of lysate.

### NanoLuc Assay in *C. elegans*

#### Growing Up and Harvesting Worms

For all NanoLuc Assays, synchronized L1s were grown at 20°C on NGM plates seeded with *E. coli* OP50-1. L1s were harvested after 4 hours (h), L2s after 20 h, L3s after 29 h, L4s after 44 h and young adults after 52 h of plating. Worms were harvested by washing off NGM plates with M9-T. If not indicated otherwise, all spins to settle worms were performed at 2,000xg for 1 minute.

#### Preparing Worm Lysate

To prepare worm lysate of 100, 1,000, 10,000 or 20,000 N2 and ERT513 young adults for Figure 1, worms were harvested and washed with M9-T. After centrifugation, the total volume including the pellet was reduced to 300 μl by removing the supernatant to the 300 μl label on the tube. Then, 300 μl of 1X Lysis Buffer with protease inhibitor (1X Lysis Buffer: 50 mM HEPES pH 7.4, 1 mM EGTA, 1 mM MgCl2, 100 mM KCl, 10% Glycerol, 0.05% NP40, 0.5 mM DTT; protease inhibitor: Millipore Sigma, cOmplete™ Cat# 11836170001) were added. After spinning down the worm pallet, the supernatant was reduced to 250 μl. The samples were then vortexed on a Disruptor Genie vortexer (Scientific Industries) with 15-20 Silicon Carbide beads in microfuge tubes for 5 minutes at 4°C. The ground samples were spun down at 20,000xg for 5 minutes and 200 μl of the worm lysate was transferred to a black, clear-bottom 96-well assay plate. NanoGlo Reagent (Promega, Nano-Glo® Luciferase Assay System, #N1110) was prepared as per manufacturer’s instructions (50 volumes NanoGlo Buffer and one volume NanoGlo Substrate) and 50 μl were added to each sample using a repeater pipette. Worm lysate and NanoGlo Reagent were briefly re-suspended by pipetting up and down with a multichannel pipette.

To prepare worm lysate of 100 and 5,000 N2, ERT513 or ERT529 worms for Figures 3, 5 and 6, animals were harvested and washed with M9-T. The worm pallet was spun down and the supernatant was removed to the 100 μl mark on the tube. 300 μl of 1X Lysis Buffer with protease inhibitor were added and after spinning the samples down the volume was reduced to 100 μl. Then, the samples were vortexed on a Disruptor Genie vortexer with ∼10-15 Silicon Carbide beads for 4 minutes at 4°C. The samples were spun down at 20,000xg for 1 minute and 50 μl of worm lysate were transferred to black, clear-bottom 96-well assay plates. To each sample, 25 μl of freshly prepared NanoGlo reagent were added.

**Figure 2.**
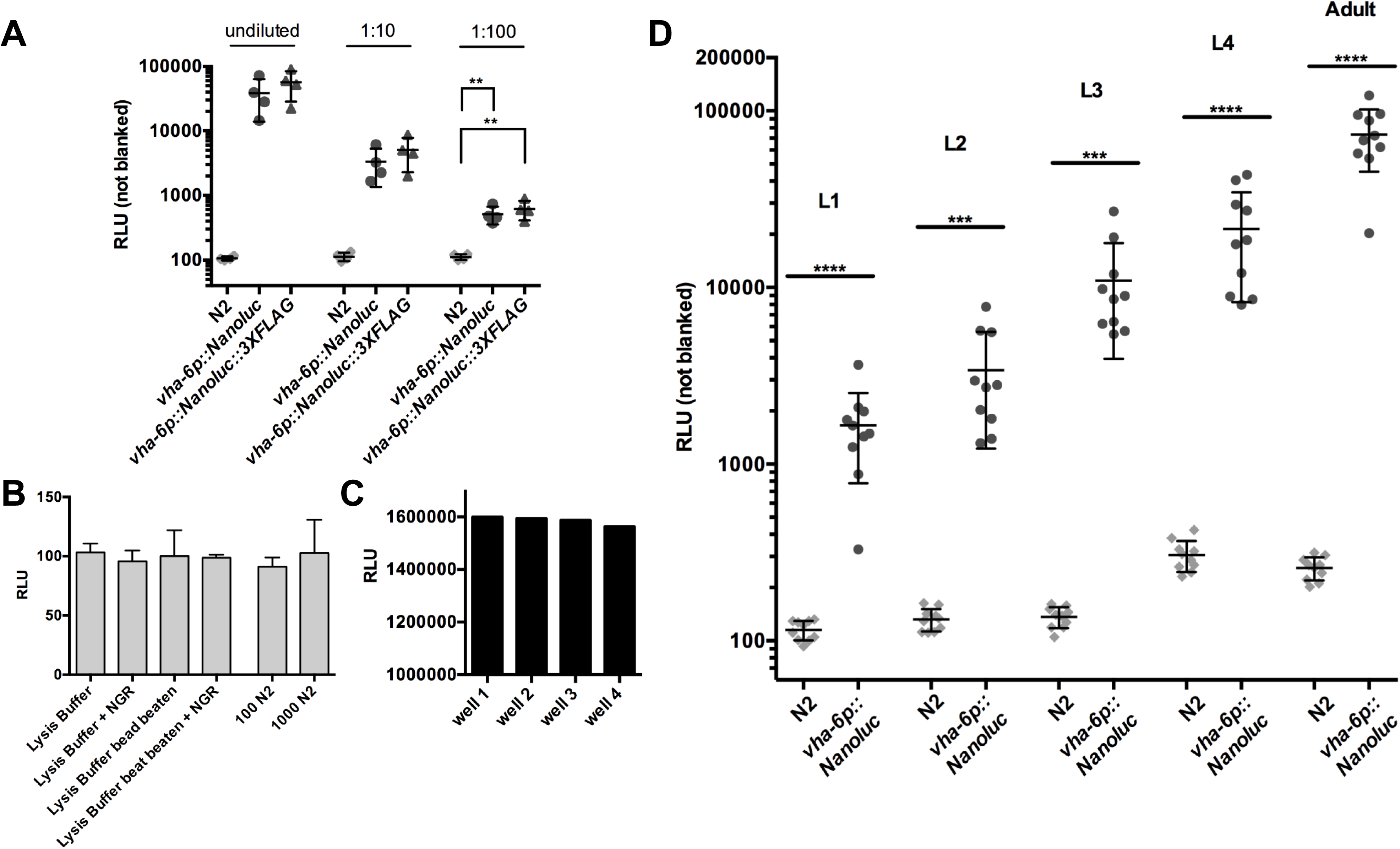
NanoLuc Signal Can Be Detected Throughout the *C. elegans* Life Cycle. (A) Bioluminescent signal measured in undiluted and diluted lysate of single *vha-6p::Nanoluc* or *vha-6p::Nanoluc::3XFLAG* adults, and N2 adults. Error bars are SD, ** *p* < 0.01. (B) Background signal of black 96-well assay plate determined with buffers, reagents and N2 lysate. Bead beating in microfuge tubes with Silicon Carbide beads performed for 5 minutes at 4°C. NGR NanoGlo Reagent, error bars are SD (n=3). (C) Signal from worm lysate aliquots from the same sample measured in multiple wells of a black 96-well assay plate (wells B10, C10, D10 and E10). (D) Each dot is the signal from a single worm. Data show robust NanoLuc activity in lysate of *vha-6p::Nanoluc* adults throughout the life cycle, but not N2 adults. Error bars are SD, **** *p* < 0.0001, *** *p* < 0.001.

**Figure 3.**
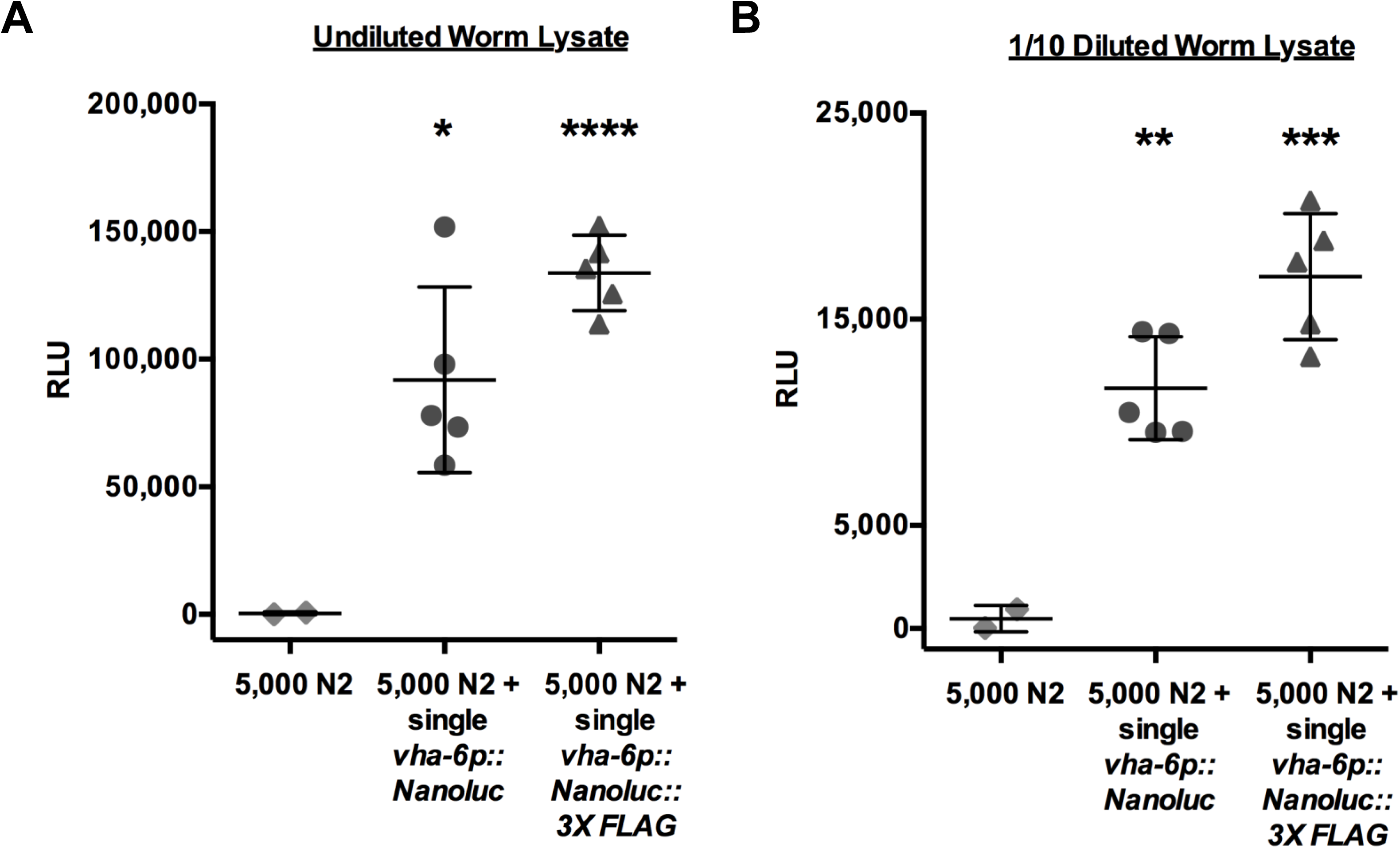
NanoLuc is a Highly Sensitive Genetic Reporter in the *C. elegans* Intestine. (A) Bioluminescent signal measured in the lysate of 5,000 N2 adults mixed with a single N2 adult, a single adult expressing *vha-6p::Nanoluc* or *vha-6p::Nanoluc::3XFLAG*. (B) Signal of 1/10 dilution of the samples measured for Figure 3A. Error bars are SD, **** *p* < 0.0001, *** *p* < 0.001, ** *p* < 0.01, * *p* < 0.05.

**Figure 4.**
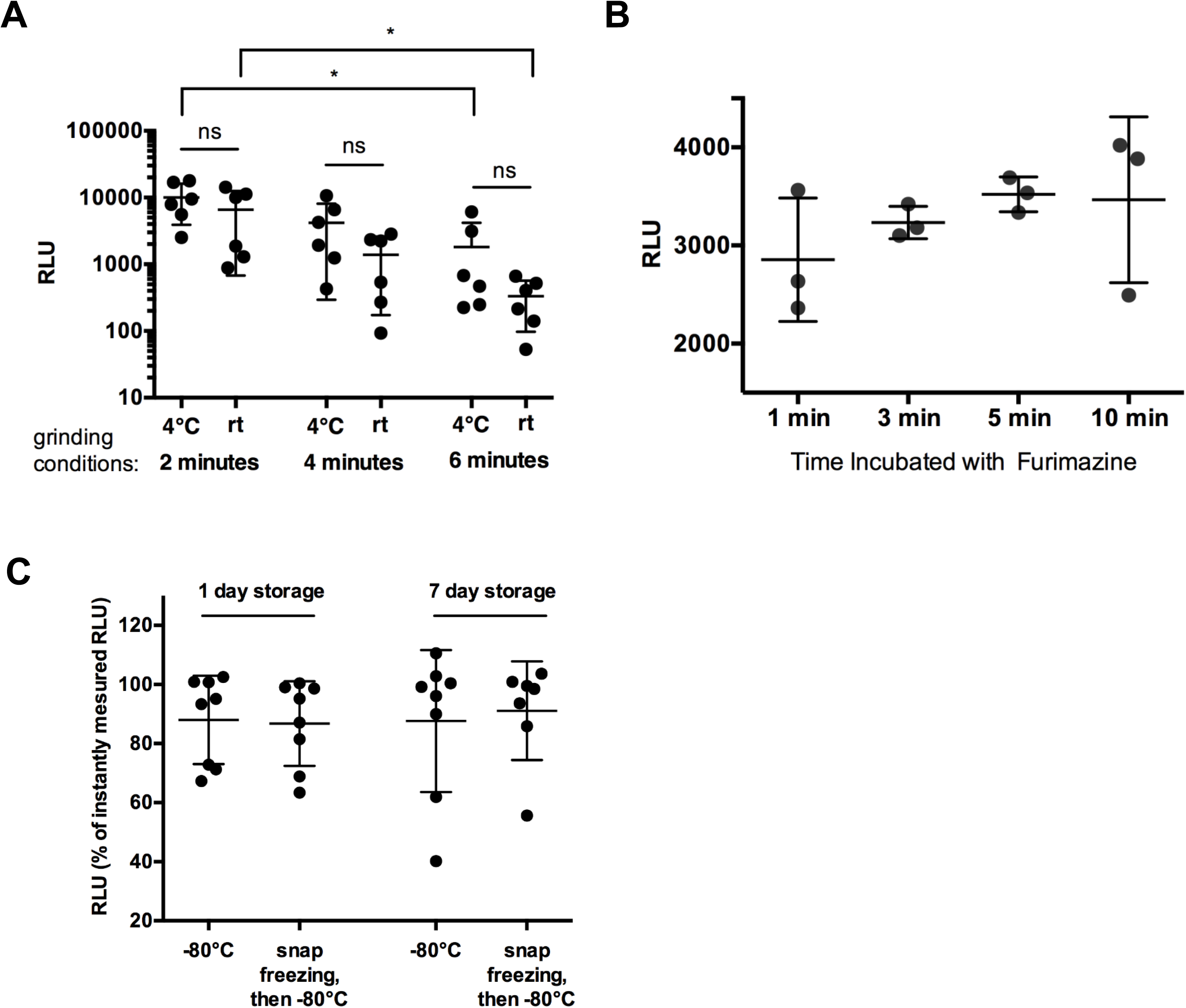
Optimizing Grinding Conditions, Substrate Incubation Time and Sample Storage for NanoLuc Assay. (A) Bioluminescent signal measured in lysate of single *vha-6p::Nanoluc* adults obtained by vortexing for two, four, and six minutes with Silicon Carbide beads at 4°C or room temperature. rt = room temperature, * *p* < 0.05, ns = not significant. (B) Bioluminescent signal measured in *vha-6p::Nanoluc* worm lysate immediately after harvesting or storing for one and seven days at - 80°C with or without initial liquid Nitrogen snap freezing. Measurements displayed as percentages of signal measured immediately after harvesting. (C) Bioluminescent signal measured in lysate of single *vha-6p::Nanoluc* adults incubated with Furimazine for one, three, five, and ten minutes. (A-C) Error bars are SD.

**Figure 5.**
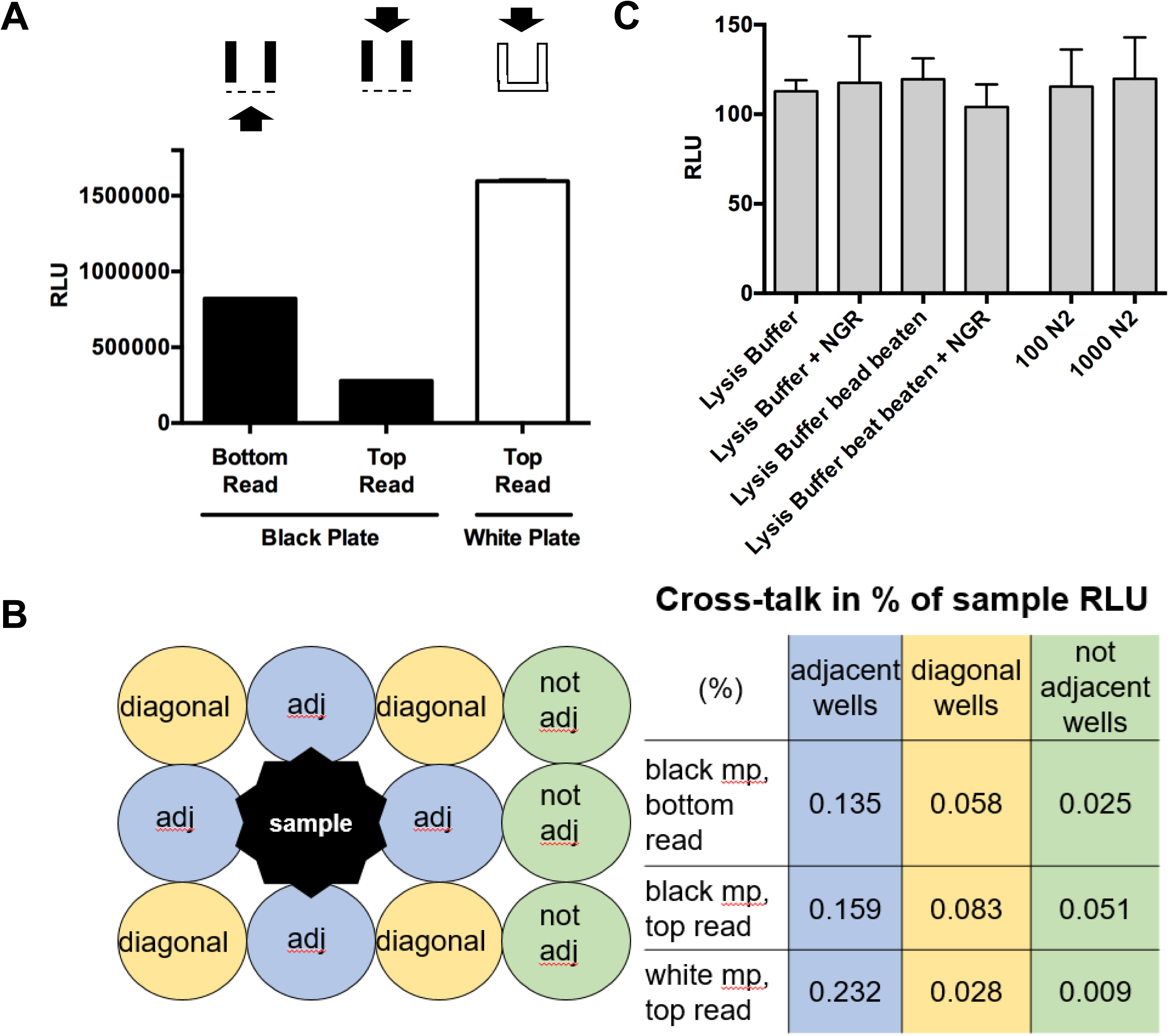
White 96-well Assay Plates Increase Detectable Signal with Minimal Cross-talk and Background Signal. (A) Bioluminescent signal from lysate aliquots of 50 *vha-6p::Nanoluc* adults measured in either black or white 96-well assay plates. Black plates had clear bottoms, white plates were completely opaque. Signal was detected from either the bottom or the top of wells as indicated. Error bars are SD (n=2) (B) Signal from lysate of 50 *vha-6p::Nanoluc* adults measured in adjacent (adj), diagonal or not adjacent (not adj) wells on 96-well assay plates. Cross-talk expressed as percentage of signal in sample well (1.5 Million RLU). mp = multi-plate. (C) Background signal of white 96-well assay plate determined with buffers, reagents and N2 lysate. Bead beating in microfuge tubes with Silicon Carbide beads performed for 5 minutes at 4°C. NGR NanoGlo Reagent, error bars are SD (n=3).

**Figure 6.**
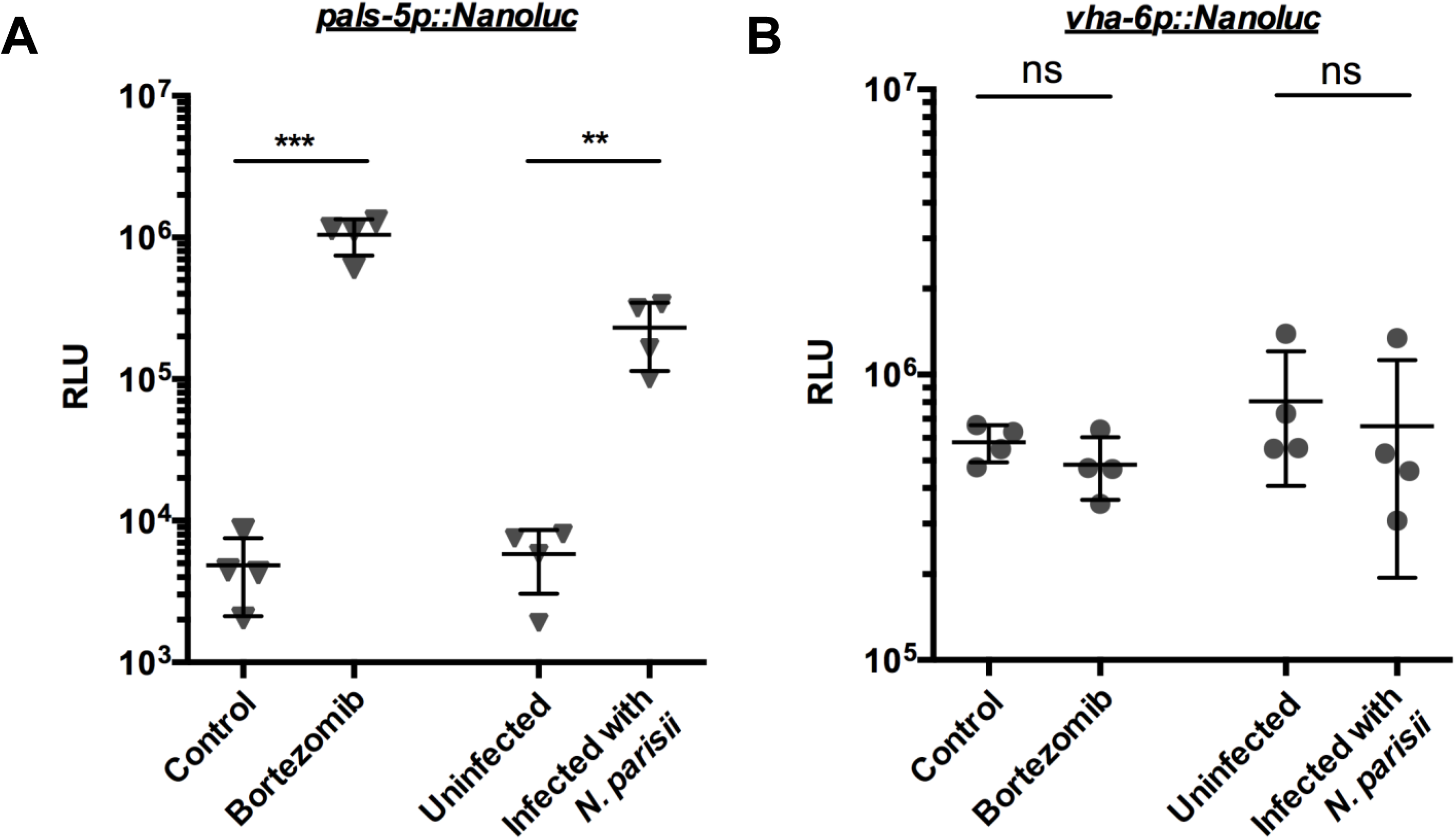
NanoLuc as an Inducible Genetic Reporter in *C. elegans*. (A,B) 100 *pals-5p::Nanoluc* (A) and *vha-6p::Nanoluc* (B) L4s treated with 22 uM Bortezomib or infected with 166,000 *N. parisii* spores, for five and four hours, respectively. Bioluminescent signal measured in mock treated vs. bortezomib treated worms or uninfected vs. *N. parisii* infected worms. One data point represents two independent measurements performed on the same day. Error bar is SD, *** *p* < 0.001, ** *p* < 0.01, ns not significant.

To prepare worm lysate of single worms for Figures 2 and 4, an eyelash pick was used to transfer individual N2, ERT513 or ERT529 worms to microfuge tubes pre-filled with 80 μl 1X Lysis Buffer. 5-10 Silicon Carbide beads were added and samples were vortexed on a Disruptor Genie vortexer for 4 minutes at 4°C. The samples were spun down at 20,000xg for 1 minute and 50 μl of worm lysate were transferred to black, clear-bottom 96-well assay plates. To prepare dilutions, wells were prefilled with appropriate amounts of 1X Lysis Buffer and worm lysate was added to a total volume of 50 μl. To each sample, 25 μl of freshly prepared NanoGlo reagent were added.

For storage experiments in Figure 4B, worm lysate was frozen to −80°C in 1X Lysis Buffer. Worm lysate samples were either placed to −80°C immediately after harvesting or first flash frozen in liquid N2 for a couple of seconds. To measure the bioluminescent signal, samples were thawed at room temperature and signal was obtained as previously described.

#### Measuring the NanoLuc Assay

The assay plate was agitated for 10 minutes at room temperature and luminescent signal was detected on a NOVOstar plate reader for 1 second without filters. If not indicated otherwise, the optimal gain across the plate was determined from the sample with the highest luminescent signal. 1X Lysis Buffer vortexed with Silicon Carbide beads and NanoGlo Reagent was used as a blank.

If not indicated otherwise, all assays were measured using black, clear-bottom 96-well assay plates (Costar, #3603) and signal was read from the bottom. Reading signal from top was tested in black plates and in opaque white plates (Costar, #3912) for Figure 5 and Supplementary Figures 1 and 2.

#### Comparing NanoLuc Signal in Intact and Lysed Worms for Supplementary Figure 1

10,000 *pals-5p::GFP* (not expressing NanoLuc, *jyIs8*) or *vha-6p::Nanoluc* (ERT513) adult worms were harvested into 15 ml tubes and washed with M9-T to remove residual bacteria. After spinning the worm pallet down, the supernatant was reduced to the 1 ml marking on the tube, and 1 ml of 2X Lysis Buffer with protease inhibitor was added to generate a worm suspension with 5 worms/μl. To measure the signal in 100 intact worms, 20 μl of worm suspension and 30 μl 2X Lysis Buffer were combined in wells of an opaque white 96-well plate. To measure the signal in 100 lysed worms, 20 μl of worm suspension, 40 ul 2X Lysis Buffer and 5 Silicon Carbide beads were combined in microfuge tubes and vortexed on a Disruptor Genie vortexer for 4 minutes at 4°C. The samples were spun down at 20,000xg for 1 minute and 50 μl of worm lysate were transferred to an opaque white 96-well assay plate. To each sample, 50 μl of freshly prepared NanoGlo reagent were added. The assay plate was agitated at room temperature and luminescent signal was detected after 10 and 60 minutes on a NOVOstar plate reader for 1 second without filters. Two measurements were performed at each time point, using either the optimal gain 2662 across all samples or the maximum gain 4095. 2X Lysis Buffer vortexed with Silicon Carbide beads and NanoGlo Reagent was used as a blank.

#### Preparing Worm Lysate Directly in 96-well Assay Plates for Supplementary Figure 2

100 *pals-5p::Nanoluc* or *vha-6p::Nanoluc* expressing adult worms were harvested into 1.5 ml tubes and washed with M9-T to remove residual bacteria, the supernatant was reduced to a volume of 100 μl. To prepare worm lysate in tubes, five Silicon Carbide beads were added and the samples were vortexed on a Disruptor Genie vortexer for 4 minutes at 4°C. Then, the samples were spun down for 1 minute at 20,000xg, and 50 μl of the supernatant were transferred to an opaque, white 96-well assay plate. To prepare worm lysate directly inside a well of an assay plate, the 100 ul worm suspension was transferred into a well of an opaque, white 96-well assay plate. Five Silicon Carbide beads were added; the plate was sealed with sealing tape (BioRad, Cat# MSB1001), fastened to a vortexer using an attachment (Scientific Industries, Cat# 504-0233-00 Model H301) and vortexed for 4 minutes at 4°C. Then, the plate was spun down at max speed for 1 minute. 50 ul of the supernatant were transferred to a fresh well. Luminescent signal was measured in lysate prepared in microfuge tubes or lysate (with or without Silicon Carbide beads) prepared in wells of an assay plate as described previously.

### Testing Inducible Genetic Reporter Induction

100 starved *vha-6p::Nanoluc* and *pals-5p::NanoLuc* L1s were grown for 48 h to the fourth larval state (L4) at 20°C. Then, the animals were treated or DMSO mock-treated with Bortezomib or infected with *N. parisii* (*ERTm1*). All biological replicates were prepared in duplicate. For bortezomib treatment, a 10 mM stock of bortezomib in DMSO was diluted with M9, then NGM plates were top plated to reach a final concentration of 22 bortezomib in the plates. For mock-treatment, DMSO and M9 were top plated, respectively. Animals were incubated for 5 h at 20°C before harvesting and analysis. For infection with *N. parisii* (ERTm1), 166,000 spores in M9 and 50 μl OP50 or M9 and 50 μl OP50 only for mock-treatment was top plated. Animals were incubated for 4 h at 20°C before harvesting and analysis. The worms were harvested and washed with M9-T and the NanoLuc Assay was performed as described in “Preparing Worm Lysate” for 100 worms and “Measuring the NanoLuc Assay”.

### Statistical Analysis of Expression Data

Statistical significance was determined with the parametric Student’s t-test for comparing two unpaired groups.

### Statement on Data Availability

Strains and plasmids are available upon request. The authors affirm that all data necessary for confirming the conclusions of the article are present within the article, figures, and tables.

## Results

### Comparisons of GFP and NanoLuc Signal in transgenic *C. elegans* adults

With the goal of developing a sensitive, plate-based assay to measure reporter gene expression in *C. elegans*, we compared signal from transgenically expressed GFP or NanoLuc under similar conditions. First, we analyzed animals that contain an integrated, single-copy 3X FLAG-tagged GFP controlled by the *vha-6* promoter. (Of note, the 3XFLAG tag serves the purpose of facilitating subsequent biochemical analysis.) The *vha-6* promoter is a commonly used promoter in *C. elegans* that drives strong, constitutive expression in the intestine (Oka et al. 2001). The strong intestinal GFP expression from these *vha-6p::GFP::3XFLAG* adult animals can be visualized with standard microscopy (Figure 1A and B). We used a conventional plate reader (NOVOstar, BMG Labtech) to measure green fluorescent signal in 100 - 20,000 intact *vha-6p::GFP::3XFLAG* adults compared to non-transgenic wild-type N2 controls. The plate reader only detected a 3 and 6-fold higher fluorescent signal in 20,000 and 10,000 *vha-6p::GFP::3XFLAG* animals compared to 20,000 and 10,000 N2 animals, respectively (Figure 1C). There was no significant signal over background when only 1,000 or 100 animals were measured. In an attempt to increase signal, we lysed worms first and then measured green fluorescent signal. Here, the plate reader did not detect any significant signal over background in lysate generated from 20,000, 10,000, 1,000 or 100 *vha-6p::GFP::3XFLAG* adult animals.

Because of the poor signal from GFP measured on a plate-reader, we explored a NanoLuc-based assay. First, we generated transgenic animals that contain a single-copy *vha-6p::Nanoluc* transgene. Using this strain, we found that luminescence from intact animals was detectable, but highly variable and did not correlate well with input amounts (Supplementary Figure 1). Therefore, we lysed worms prior to performing the assay, to facilitate access of substrate to the NanoLuc enzyme and improve the signal. In contrast to the measurements from intact animals, we detected a robust and reproducible signal from lysates of NanoLuc-expressing animals. Impressively, we detected a 400,000-fold higher signal in lysate from 100 *vha-6p::Nanoluc* adult animals when compared to lysate from 100 N2 animals (Figure 1C). Lysates from 1,000 *vha-6p::Nanoluc* adult animals had significantly higher signal, and lysates from 10,000 and 20,000 *vha-6p::Nanoluc* adult animals saturated the detection capability of the plate reader (Figure 1C). Because of the higher levels of bioluminescent signal obtained from lysates compared to intact animals, we used lysates for all subsequent analyses (see Figure 7 for Assay Schematic). Overall, these results demonstrate a robust and reproducible signal in a plate reader-based assay using lysate from *C. elegans* expressing NanoLuc, which is much greater than the signal from GFP under similar conditions.

**Figure 7.**
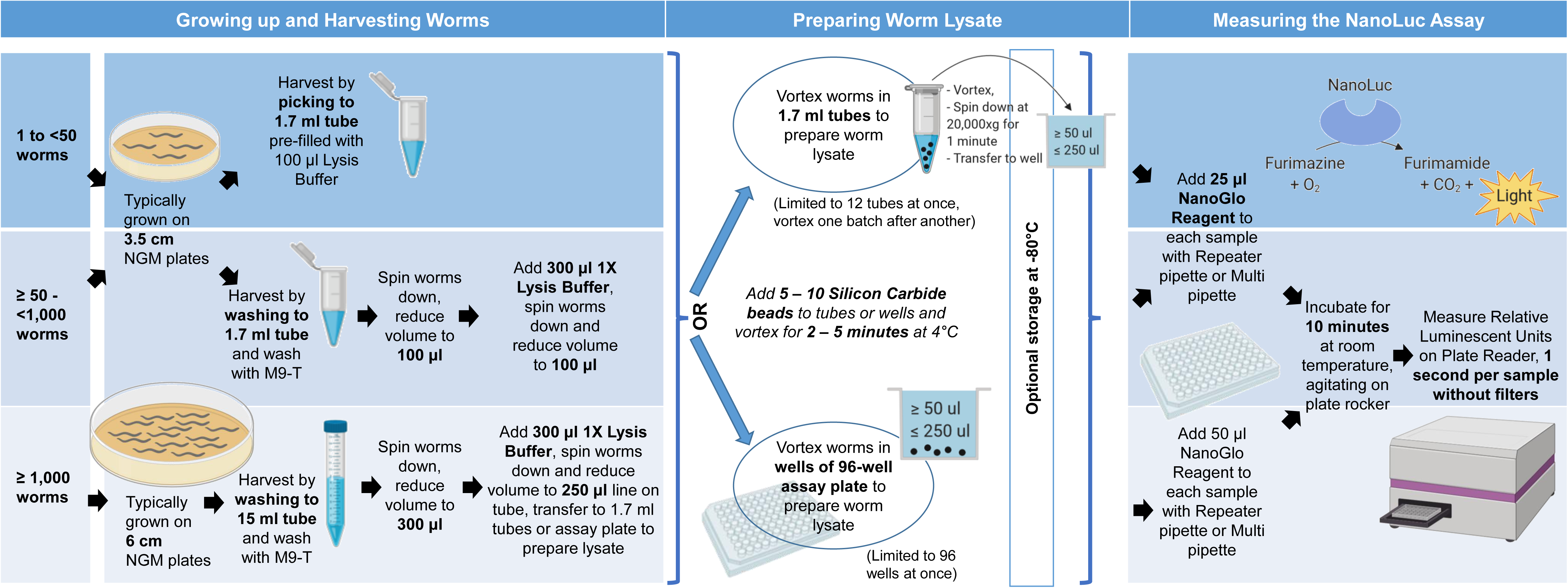
Schematic of the *C. elegans* NanoLuc Assay Workflow. The amount of worms needed for an assay depends on the strength of expression from the promoter in use and can be measured in single animals or in populations of tens of thousands of animals. We routinely use 100 L4s to detect activation of a *pals-5* stress/immune pathway. Worms are single picked to 1.7 ml tubes pre-filled with Lysis Buffer, or washed to 1.7 ml or 15 ml tubes with M9 buffer with 0.1% Tween 20 (M9-T) and subsequently washed with M9-T to remove residual bacteria. The main goal is to harvest and lyse the worms in as little volume as possible. After spinning the worms down, the supernatant is removed and then washed with 1X Lysis Buffer containing protease inhibitor. Lysate can be prepared either inside 1.7 ml tubes or directly in wells of a white, opaque 96-well assay plate. Worm lysate is prepared by adding 5 – 10 Silicon Carbide beads to the worm suspension in tubes or wells, then closing tubes or sealing the wells and vortexing for 4 minutes at 4°C. Optionally, worm lysate can be stored for several days at −80°C after this step. The tubes or the assay plate are/is spun down at 20,000xg for 1 minute. If grinding in tubes, at least 50 µl of worm lysate are transferred to wells of white, opaque 96-well assay plate. 25 – 50 µl of NanoGlo Reagent (Promega) are added to worm lysate and the plate is agitated for 10 minutes at room temperature on plate rocker. With a plate reader, Relative Luminescent Units are measured by detecting luminescent signal for 1 second per sample without filters. As a blank, Lysis Buffer is vortexed with Silicon Carbide beads and NanoGlo Reagent is added 10 minutes before measurement. The assay gain is set so that the highest signal does not saturate the detector capability of the photo multiplier tube. PI = protease inhibitor, µl = microliter.

### Assessing the Sensitivity of NanoLuc Constitutively Expressed in the Intestine

To assess the sensitivity of NanoLuc signal in *C. elegans*, we measured the reporter signal in lysate from single *vha-6p::Nanoluc* expressing animals at different life stages (Figure 2). We detected a robust luminescent signal in single adults that was significantly higher than the N2 background (Figure 2A). We also generated a *vha-6p::Nanoluc::3XFLAG* strain and found similar results as the *vha-6p::Nanoluc* strain, with robust signal detected in single adults (Figure 2A). Further, we detected strong signal in 10- and 100-fold dilutions of lysates made from single adult *vha-6p::Nanoluc* and *vha-6p::Nanoluc::3XFLAG* animals (Figure 2A). The background signal of Lysis Buffer, substrate and lysed N2 was similar to the plate reader background determined with water, indicating very low background for this assay (Figure 2B). Importantly, we saw a very stable signal throughout different wells (B10, C10, D10, E10) of a 96-well assay plate (Figure 2C). To assess NanoLuc signal sensitivity in younger animals, we measured the signal in all four larval stages (L1 through L4) and in young adults. We detected a robust signal in single worms from all of these life stages (Figure 2D). In summary, a significant signal over background can be detected throughout all *C. elegans* life stages from single animals constitutively expressing NanoLuc in the intestine (Figure 2D). Further, signal can be detected in as little as a 1/100 dilution of lysate from a single adult animal (Figure 2A).

Besides detecting low reporter expression, detecting rare events can be critical to the success of reporter-based assays. Therefore, we mimicked a rare event by mixing a single NanoLuc-expressing worm in a pool of 5,000 N2s. Here, we detected an approximately 100,000-fold and 12,000-fold signal over N2 background in undiluted worm lysate and 1:10 diluted worm lysate, respectively (Figure 3A and B). Therefore, this NanoLuc assay should allow detection of reporter expression coming from only a single animal in a large population of non-expressing animals.

### Optimizing Lysate Preparation Conditions for NanoLuc Assay

Next, we tested several assay parameters for NanoLuc detection, including grinding conditions, substrate incubation time and sample storage. The worm lysate described for the NanoLuc assays above was generated by vortexing worms in microfuge tubes with Silicon Carbide beads on a vortexer. We compared grinding times of two, four, and six minutes at 4°C or at room temperature. We found that signal is decreased with longer grinding times (Figure 4A). There was no significant difference between grinding at 4°C and at room temperature, although there was a trend toward lower signal at room temperature. In cell culture studies, bioluminescent NanoLuc signal is usually measured after three minutes of incubation of cell lysate with Furimazine. Our data show that worm lysate can be incubated with substrate for three to ten minutes before measuring (Figure 4B).

The workflow of many experiments can benefit from the option to store samples for several days prior to data collection. To evaluate if NanoLuc samples can be stored before measurement, we measured signal in *vha-6p::Nanoluc* worm lysate immediately after worm lysis, after one day of storage at −80°C, and after seven days of storage. We either froze down the lysate directly or after snap freezing in liquid nitrogen. With both storage methods we were able to preserve the bioluminescent signal for up to seven days without significant signal loss (Figure 4C). In summary, our analyses indicate that worm lysate should be prepared by grinding the worms for two to four minutes, then measured immediately or stored at −80°C for several days. The lysate can be incubated with substrate for up to ten minutes before assessing the bioluminescent signal.

### Testing Multi-well Plate Assay Parameters for NanoLuc Assay

Next, we optimized plate-reading parameters. While black assay plates are recommended for detecting fluorescence because they minimize background, white assay plates are recommended for bioluminescence because of the increased signal possible due to the reflection that occurs in white plates together with the low background typical for bioluminescent assays (Judy Gibbs 2001; Wohlstadter et al. 2005). Another assay parameter to consider is whether the signal is detected from the top or the bottom of the plate. Top reading has been reported to minimize well-to-well cross-talk compared to bottom reading because the detector optics can function as a lid for the sample well and shield it from incoming light (Bjerke 2014). Therefore, we compared luminescent signals measured from the top of opaque white assay plates to bottom or top reading of the previously used black, clear bottom assay plates. We detected a twice higher signal in aliquots of the same sample when we measured in white plates from the top compared to measuring in black plates from the bottom (Figure 5A). This effect was not compensated by measuring in black plates from the top, as the signal is even four-fold lower than bottom reading in the same plates. Of note, both black and white plates show minimal well-to-well crosstalk below 1% of sample signal in wells adjacent and diagonal from a sample with strong signal (1.5 Million RLU, Figure 5B). These results indicate that white plates increase the detectable bioluminescent signal in *C. elegans* without increasing background (Figure 5C) and maintaining minimal cross-talk.

### Demonstration of NanoLuc as a Sensitive Reporter for Monitoring Inducible Gene Expression

Finally, we investigated whether NanoLuc could be used as an inducible genetic reporter. Here we generated animals with a single-copy NanoLuc driven by the *pals-5* promoter. The *pals-5* gene is used as a read-out for the Intracellular Pathogen Response (IPR), which is a defense program induced by diverse, natural intracellular pathogens, as well as by proteotoxic stress (Reddy et al. 2019). Previous studies have used a multi-copy *pals-5::GFP* reporter strain to monitor *pals-5* induction in the intestine upon blockade of the proteasome, or with intracellular infection by the microsporidian *N. parisii* (Bakowski et al. 2014). With the integrated, single-copy *pals-5p::Nanoluc* transgene, we detected a ∼300-fold increase upon treatment with proteasome inhibitor bortezomib and a ∼50-fold increase in luminescent signal after infection with *N. parisii* in 100 L4 worms (Figure 6A). We controlled for changes in overall worm mass and treatment interaction with NanoLuc by measuring expression of the constitutively expressed *vha-6p::Nanoluc* in parallel. The luminescent signal in treated and untreated *vha-6p::Nanoluc* animals was not significantly different upon treatment (Figure 6B), indicating that bortezomib treatment and microsporidia infection did not simply increase NanoLuc signal independent of the *pals-5* promoter.

In order to streamline the NanoLuc assay workflow, we also tested whether signal could be measured directly from worm lysate prepared in a multi-well plate, to eliminate the need for a separate step where worms are first disrupted in microfuge tubes and then transferred into wells of an assay plate. Here, we tested two conditions: 1) we vortexed worms with Silicon Carbide beads in the wells and transferred half of this lysate away from the beads into fresh wells where it was measured, and 2) we directly measured signal from wells that contain both worm lysate and beads (Supplementary Figure 2). In Condition #1 we measured the same signal as from lysate that was prepared in microfuge tubes and transferred into wells to be measured. For Condition #2 we found that the absolute signal we measured was lower when beads were still present in the wells, but the fold-increase upon bortezomib treatment was identical to that seen for measurements of lysate only. Therefore, in cases where there is strong induction such as with the *pals-5p::NanoLuc* reporter, the signal can conveniently be measured from worms disrupted in the wells and subsequently measured from the same wells. Overall, these results indicate that NanoLuc can be used effectively both as a constitutive as well as an inducible genetic reporter in *C. elegans*.

## Discussion

Here, we describe a highly sensitive luminescence-based method to detect gene expression in *C. elegans*, using the luciferase NanoLuc. In a plate reader setting where intestinally expressed GFP could be detected only 6-fold over background levels, we found that intestinally expressed NanoLuc could be detected at several million-fold over background (Figure 1C). We used this strain that expresses NanoLuc intestinally to detect signal in lysate of single worms at all life stages from L1s to adults. The signal was so robust that we could detect it confidently over background in worm lysate dilutions as small as 1/100 of a single NanoLuc-expressing adult animal. The optimization and testing we performed of various assay conditions provide guidelines for grinding time and temperature, sample storage, and plate type. We also found that the assay can be further streamlined by grinding and measuring directly in a multi-well plate (see Schematic in Figure 7).

Our initial motivation to translate this extremely bright genetic reporter from cell culture to the nematode *C. elegans* was to monitor rare transformation events in obligate intracellular pathogens of the *C. elegans* intestine from the Microsporidia phylum (Reinke and Troemel 2015). To-date, there has been no successful genetic modification of any of the >1,400 species in the Microsporidia phylum, despite their widespread significance to the fields of agriculture, evolution and pathogenesis (Munita and Arias 2016; Vávra and Lukeš 2013). The small size, high physical stability, and high brightness of NanoLuc should enable the detection of rare, single microsporidia transformation events in the intestine of individual *C. elegans* animals within a large population being tested. Indeed, our analysis indicated that it was possible to easily detect a single NanoLuc expressing worm in a population of thousands of non-NanoLuc-expressing worms.

In addition to measuring constitutive expression, we demonstrated that NanoLuc could be used as a convenient tool to monitor inducible promoter activity, showing a 300 and 50-fold increase in expression from the *pals-5* stress/immune reporter after proteasome blockade and infection with microsporidia, respectively. To assess if those treatments influence NanoLuc activity or animal size, we used the *vha-6* promoter-driven NanoLuc expression in treated and untreated animals as a control. In the future, Dual Luciferase assays that measure promoter-of-interest::Nanoluc and control-promoter::Firefly luciferase expression in the same animal could provide an internal control (Promega, Nano-Glo® Dual-Luciferase® Reporter Assay System, #N1610). However, an advantage of a NanoLuc-only system for monitoring gene expression is that it is not affected by changes in ATP levels, which regulate luminescence from firefly and other luciferases.

The advantages and disadvantages of a NanoLuc-based assay compared to other gene expression methods are summarized in Table 2. When compared to GFP, NanoLuc provides a much more sensitive and rapid method for plate-reader-based quantitation of gene expression. However, a major disadvantage of the NanoLuc assay is that it requires lysing the worms to generate a robust signal. This step is problematic for experiments where tissue-specific information and/or sample recovery is desirable. While it is not clear why lysis is required, a likely explanation is that the substrate Furimazine cannot sufficiently penetrate the cuticle and tissues of *C. elegans*.

**Table 2.** Comparisons between NanoLuc and other Methods for measuring Gene expression.

|  | NanoLuc | GFP | qRT-PCR | smFISH |
| --- | --- | --- | --- | --- |
| **Sensitivity** | High | Low | High | High |
| **Time to unbiased quantitation** | Minutes | Hours (depending on sample size) | Hours | Hours to Days |
| **Throughput** | High | Medium | Medium | Low |
| **Requires lysis (loses tissue information)** | Yes | No | Yes | No |
| **Input material required** | Single worms | Single worms | 100's to 1000's | Single worms |
| **Read-out** | Promoter activity | Promoter activity | Endogenous mRNA | Endogenous mRNA |
| **Strain development** | Required | Required | Not required | Not required |

Although it can be desirable to measure reporter gene expression in intact worms, the lysing procedure we describe only takes a few minutes, and thus is much faster than other end-point assays for gene expression that lose tissue information, like qRT-PCR (Table 2). One excellent application for *C. elegans* expression of NanoLuc would be genetic screens where it is not necessary to recover live animals, such as in RNAi-based screens. For these screens, NanoLuc would provide better sensitivity and quantitation than GFP, and would be more scalable than qRT-PCR. In addition, the NanoLuc assay requires less input material than assays like qRT-PCR, and in some cases it is preferable to determine promoter activity instead of overall mRNA levels (Table 2). This feature highlights the application we have demonstrated in Figure 6, where a *pals-5* promoter-driven NanoLuc strain provides a reporter for induction of the IPR stress/immune pathway. This use is analogous to luciferase-based methods for measuring activation of stress and immune pathways in mammalian cells (Delhove et al. 2017), and may be generally useful for monitoring activation of many different kinds of transcriptional responses in *C. elegans*.

## Acknowledgements

We acknowledge Robert Luallen for generating strain ERT412, and Kirthi Reddy for help in generating strain ERT729. We also thank Spencer Gang, Vladimir Lazetic, Robert Luallen and Crystal Chhan for comments on the manuscript. We thank Michael David’s lab for maintenance of the NOVOstar plate reader. This work was supported by NIH under R01 AG052622 GM114139 to ERT.

## Figure Legends

**Supplementary Figure 1.**
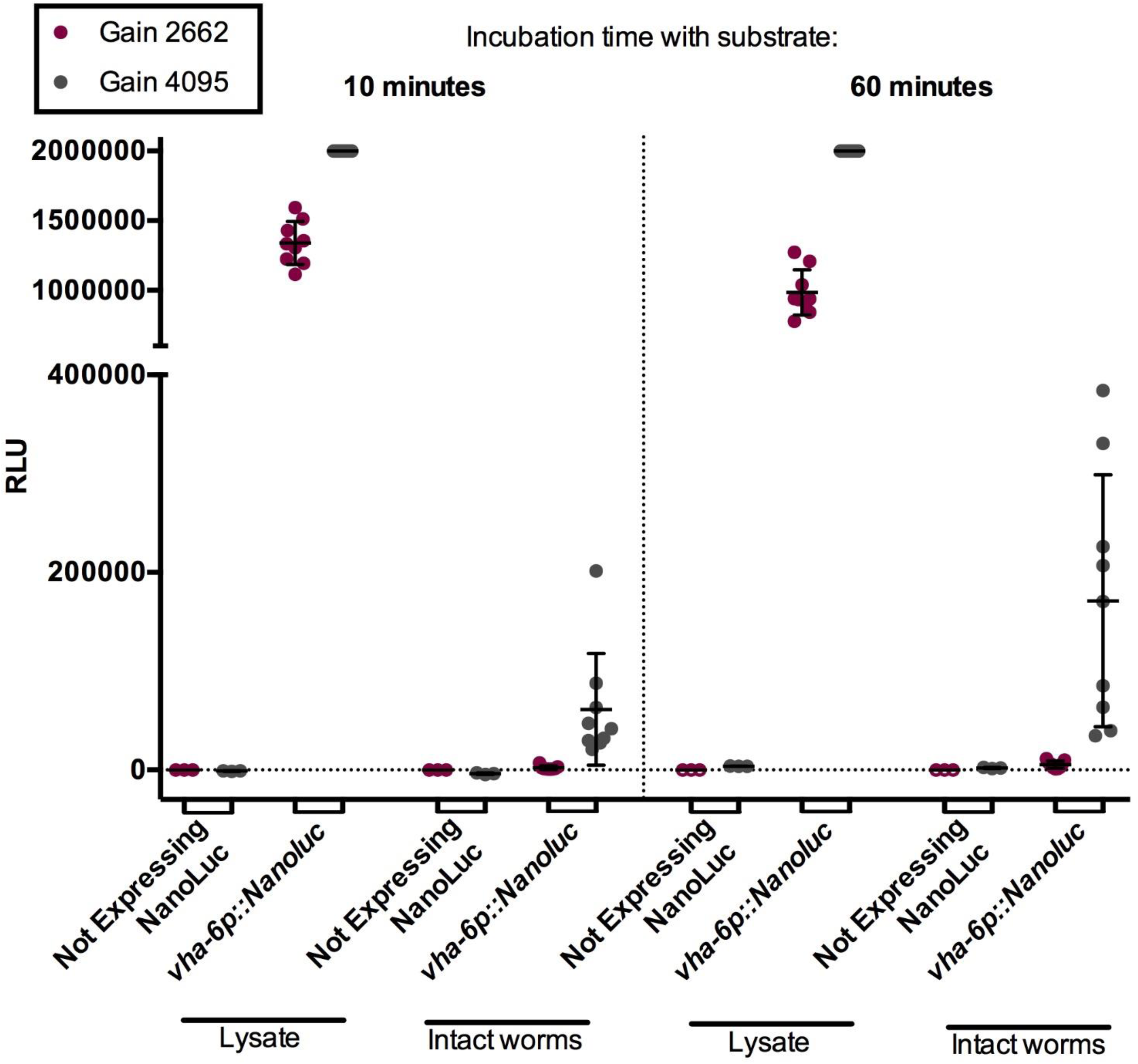
Bioluminescent Signal is Stronger and More Consistent in Lysed Worms than in Intact Worms. Bioluminescent signal measured in 100 adult worms that were intact or vortexed with Silicon Carbide beads to obtain worm lysate. Controls did not express NanoLuc but instead expressed a *pals-5p::GFP* transgene. NanoLuc expressing worms had a single copy insertion of a *vha-6p::Nanoluc* construct. Signal was measured 10 and 60 minutes after addition of the substrate Furimazine via NanoGlo Reagent (Promega). Red dots represent measurements with an optimal gain across all samples (gain 2662) and grey dots represent measurements with the maximal possible gain to determine the highest detectable luminescent signal for intact worms (max. gain 4095). With a gain of 4095, the signal from worm lysate of *vha-6p::Nanoluc* worms over-saturated the detection capability of the photomultiplier tube.

**Supplementary Figure 2.**
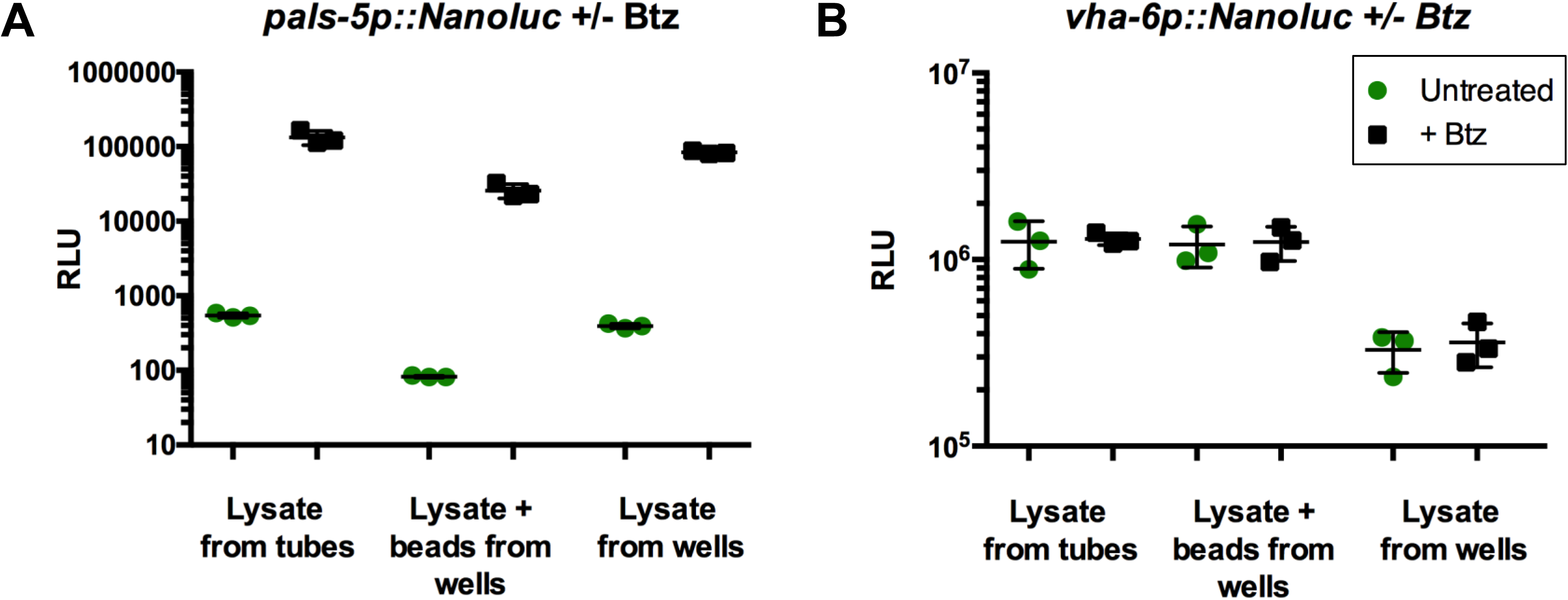
Worm Lysate Can Be Prepared and Measured Directly in Wells of 96-well Plate. (A, B) Bioluminescent signal measured in worm lysate of (A) *pals-5p::Nanoluc* or (B) *vha-6p::Nanoluc* expressing adults. Worm lysate was either prepared in 1.5 ml microfuge tubes or in wells of a 96-well assay plate. For “Lysate from tubes”, 100 ul lysate was prepared in microfuge tubes by vortexing worms, Lysis Buffer and Silicon Carbide beads. Then 50 ul of supernatant was transferred to 96-well assay plates and signal was measured. Additionally, 100 ul of lysate was prepared directly in a well of a 96-well assay plate by vortexing worms, Lysis Buffer and Silicon Carbide beads in a plate sealed with sealing tape. 50 ul of supernatant were transferred to fresh well (“Lysate from wells”) and the remaining lysate with beads was measured in presence of Silicon Carbide beads in grinding well (“Lysate + beads from wells”). Btz = bortezomib.

## Literature

Bakowski, Malina A., Christopher A. Desjardins, Margery G. Smelkinson, Tiffany L. Dunbar, Tiffany A. Dunbar, Isaac F. Lopez-Moyado, Scott A. Rifkin, Christina A. Cuomo, and Emily R. Troemel. 2014. “Ubiquitin-Mediated Response to Microsporidia and Virus Infection in C. Elegans.” PLoS Pathogens 10(6):e1004200.

Bjerke, Michael. 2014. Getting the Most from Your Plate-Based Assays.

Brenner, S. 1974. “The Genetics of Caenorhabditis Elegans.” Genetics 77(1):71–94.

Brock, Matthias. 2012. “Application of Bioluminescence Imaging for in Vivo Monitoring of Fungal Infections.” International Journal of Microbiology 2012:956794.

De-Souza, Evandro A., Henrique Camara, Willian G. Salgueiro, Raíssa P. Moro, Thiago L. Knittel, Guilherme Tonon, Silas Pinto, Ana Paula F. Pinca, Adam Antebi, Amy E. Pasquinelli, Katlin B. Massirer, and Marcelo A. Mori. 2019. “RNA Interference May Result in Unexpected Phenotypes in Caenorhabditis Elegans.” Nucleic Acids Research 47(8):3957–69.

Delhove, Juliette M. K. M., Rajvinder Karda, Kate E. Hawkins, Lorna M. FitzPatrick, Simon N. Waddington, and Tristan R. McKay. 2017. “Bioluminescence Monitoring of Promoter Activity In Vitro and In Vivo.” Methods in Molecular Biology (Clifton, N.J.) 1651:49–64.

Dokshin, Gregoriy A., Krishna S. Ghanta, Katherine M. Piscopo, and Craig C. Mello. 2018. “Robust Genome Editing with Short Single-Stranded and Long, Partially Single-Stranded DNA Donors in Caenorhabditis Elegans.” Genetics 210(3):781–87.

Emmons, S. W., M. R. Klass, and D. Hirsh. 1979. “Analysis of the Constancy of DNA Sequences during Development and Evolution of the Nematode Caenorhabditis Elegans.” Proceedings of the National Academy of Sciences of the United States of America 76(3):1333–37.

England, Christopher G., Emily B. Ehlerding, and Weibo Cai. 2016. “NanoLuc: A Small Luciferase Is Brightening Up the Field of Bioluminescence.” Bioconjugate Chemistry 27(5):1175–87.

Frøkjaer-Jensen, Christian, M. Wayne Davis, Christopher E. Hopkins, Blake J. Newman, Jason M. Thummel, Søren-Peter Olesen, Morten Grunnet, and Erik M. Jorgensen. 2008. “Single-Copy Insertion of Transgenes in Caenorhabditis Elegans.” Nature Genetics 40(11):1375–83.

Gammon, Don B., Takao Ishidate, Lichao Li, Weifeng Gu, Neal Silverman, and Craig C. Mello. 2017. “The Antiviral RNA Interference Response Provides Resistance to Lethal Arbovirus Infection and Vertical Transmission in Caenorhabditis Elegans.” Current Biology: CB 27(6):795–806.

Hall, Mary P., James Unch, Brock F. Binkowski, Michael P. Valley, Braeden L. Butler, Monika G. Wood, Paul Otto, Kristopher Zimmerman, Gediminas Vidugiris, Thomas Machleidt, Matthew B. Robers, Hélène A. Benink, Christopher T. Eggers, Michael R. Slater, Poncho L. Meisenheimer, Dieter H. Klaubert, Frank Fan, Lance P. Encell, and Keith V. Wood. 2012. “Engineered Luciferase from a Deep Sea Shrimp a Novel Imidazopyrazinone Substrate.” ACS Chemical Biology 7(11):1848.

Judy Gibbs. 2001. “Selecting the Detection System - Colorimetric, Fluorescent, Luminescent Methods ELISA Technical Bulletin - No. 5.” Corning Incorporated Life Sciences, 14.

Lagido, Cristina, Debbie McLaggan, and L. Anne Glover. 2015. “A Screenable In Vivo Assay for Mitochondrial Modulators Using Transgenic Bioluminescent Caenorhabditis Elegans.” Journal of Visualized Experiments: JoVE (105):e53083.

Mello, C. C., J. M. Kramer, D. Stinchcomb, and V. Ambros. 1991. “Efficient Gene Transfer in C.Elegans: Extrachromosomal Maintenance and Integration of Transforming Sequences.” The EMBO Journal 10(12):3959–70.

Mendenhall, Alexander R., Patricia M. Tedesco, Bryan Sands, Thomas E. Johnson, and Roger Brent. 2015. “Single Cell Quantification of Reporter Gene Expression in Live Adult Caenorhabditis Elegans Reveals Reproducible Cell-Specific Expression Patterns and Underlying Biological Variation.” PloS One 10(5):e0124289.

Minkina, Olga and Craig P. Hunter. 2018. “Intergenerational Transmission of Gene Regulatory Information in Caenorhabditis Elegans.” Trends in Genetics: TIG 34(1):54–64.

Munita, Jose M. and Cesar A. Arias. 2016. “Mechanisms of Antibiotic Resistance.” Microbiology Spectrum 4(2).

Oka, T., T. Toyomura, K. Honjo, Y. Wada, and M. Futai. 2001. “Four Subunit a Isoforms of Caenorhabditis Elegans Vacuolar H+-ATPase. Cell-Specific Expression during Development.” The Journal of Biological Chemistry 276(35):33079–85.

Olmedo, María, Mirjam Geibel, Marta Artal-Sanz, and Martha Merrow. 2015. “A High-Throughput Method for the Analysis of Larval Developmental Phenotypes in Caenorhabditis Elegans.” Genetics 201(2):443–48.

Palikaras, Konstantinos and Nektarios Tavernarakis. 2016. “Intracellular Assessment of ATP Levels in Caenorhabditis Elegans.” BIO-PROTOCOL 6(23).

Reddy, Kirthi C., Tal Dror, Ryan S. Underwood, Guled A. Osman, Corrina R. Elder, Christopher A. Desjardins, Christina A. Cuomo, Michalis Barkoulas, and Emily R. Troemel. 2019. “Antagonistic Paralogs Control a Switch between Growth and Pathogen Resistance in C. Elegans” edited by J. J. Collins. PLOS Pathogens 15(1):e1007528.

Reinke, Aaron W. and Emily R. Troemel. 2015. “The Development of Genetic Modification Techniques in Intracellular Parasites and Potential Applications to Microsporidia.” PLoS Pathogens 11(12):e1005283.

Schweinsberg, Peter J. and Barth D. Grant. 2013. “C. Elegans Gene Transformation by Microparticle Bombardment.” WormBook: The Online Review of C. Elegans Biology 1.

Teuscher, Alina C. and Collin Y. Ewald. 2018. “Overcoming Autofluorescence to Assess GFP Expression During Normal Physiology and Aging in Caenorhabditis Elegans.” Bio-Protocol 8(14).

Thorne, Natasha, James Inglese, and Douglas S. Auld. 2010. “Illuminating Insights into Firefly Luciferase and Other Bioluminescent Reporters Used in Chemical Biology.” Chemistry & Biology 17(6):646–57.

Vávra, Jiří and Julius Lukeš. 2013. “Microsporidia and ‘The Art of Living Together.’” Advances in Parasitology 82:253–319.

Wohlstadter, Jacob N., Eli Glezer, James Wilbur, George Sigal, Kent Johnson, Charles Clinton, Alan Kishbaugh, Bandele Jeffrey-Coker, Jeff D. Debad, and Alan B. Fisher. 2005. “Assay Plates, Reader Systems And Methods For Luminescence Test Measurements.” 2(60).

